# Reward certainty and preference bias selectively shape voluntary decisions

**DOI:** 10.1101/832311

**Authors:** Wojciech Zajkowski, Dominik Krzemiński, Jacopo Barone, Lisa Evans, Jiaxiang Zhang

## Abstract

Choosing between equally valued options can be a conundrum, for which classical decision theories predicted a prolonged response time (RT). Paradoxically, a rational decision-maker would need no deliberative thinking in this scenario, as outcomes of alternatives are indifferent. How individuals choose between equal options remain unclear. Here, we characterized the neurocognitive processes underlying such voluntary decisions, by integrating advanced cognitive modelling and EEG recording in a probabilistic reward task, in which human participants chose between pairs of cues associated with identical reward probabilities at different levels. We showed that higher reward certainty accelerated RT. At each certainty level, participants preferred to choose one cue faster and more frequently over the other. The behavioral effects on RT persisted in simple reactions to reward cues. By using hierarchical Bayesian parameter estimation for an accumulator model, we showed that the certainty and preference effects were independently associated with the rate of evidence accumulation during decisions, but not with visual encoding or motor execution latencies. Time-resolved multivariate pattern classification of EEG evoked response identified significant representations of reward certainty and preference choices as early as 120 ms after stimulus onset, with spatial relevance patterns maximal in middle central and parietal electrodes. Furthermore, EEG-informed computational modelling showed that the rate of change between N100 and P300 event-related potentials reflected changes in the model-derived rate of evidence accumulation on a trial-by-trial basis. Our findings suggested that reward certainty and preference collectively shaped voluntary decisions between equal options, providing a mechanism to prevent indecision or random behavior.

## Introduction

Cognitive flexibility enables decision strategies adaptive to environmental and motivational needs (Schiebener and Brand, 2015). One characteristic of this ability is that harder decisions often take longer. Evidence from neurophysiology (Gold and Shadlen, 2001), neuroimaging (Heekeren et al., 2008) and modelling (Smith and Ratcliff, 2004) suggested an evidence accumulation process for decision-making: information is accumulated over time, and a decision is made when the accumulated evidence reached a threshold (Gold and Shadlen, 2007). According to this framework, decision difficulty, and in turn response time (RT), is proportional to the relative difference in the evidence supporting each option, consistent with results from perceptual (Ditterich et al., 2003) and value-based (Oud et al., 2016) decisions.

What happens if decision difficulty reaches a tipping point with values of options being equal? Classical stochastic decision models would predict a deadlock or indecision between equal choices (Pais et al., 2013). In reality, individuals can choose between options with indistinguishable certainty of receiving rewards (Voigt et al., 2019), such as deciding between well-liked dishes in a good restaurant, or between underwhelming takeaways for a casual meal. Paradoxically, economic analysis suggests that the closer options are in their reward probability, the less beneficial to wait longer to decide. The benefit of “rushing to decisions” increases as reward certainty gets larger because cognitive resources could be relocated elsewhere (Rustichini, 2009), predicting faster responses with higher reward certainty (Pirrone et al., 2018). Furthermore, if choices are purely based on expected rewards, one may choose equal-valued options randomly with the same probability. Nevertheless, choice behaviour deviates from random in both lab-based (Zhang and Rowe, 2015) and consumer decisions (Wheeler, 1974), suggesting a possible bias between equal options, rendering some options more preferred than others.

Previous research on equal choices raises three further unresolved issues. First, it is unclear how reward certainty and preference bias impact on different sub-components of the decision-making process. Second, functional imaging studies have localized the mesocorticolimbic dopaminergic network to be involved in both reward certainty and preference processing (Tobler et al., 2007; Abler et al., 2009). Less is known about how macroscopic brain activities relating to these effects unfold in time. Third, conventional equal-choice paradigms commonly use subjective ratings to quantify and equate values of options. This design has been shown to be vulnerable to value fluctuations (Chen and Risen, 2010; Izuma and Murayama, 2013) that originate from voluntary choices themselves (Festinger, 1957; Bem, 1967).

Here, we addressed these questions by combining advanced computational modelling and EEG in a probabilistic reward task. Participants memorized six unambiguous cues associated with three levels of reward certainty, and they made binary choices between cues with equal reward certainty. Additional control conditions involved decisions between cues with unequal certainty and unitary responses to single cues. This design enabled us to focus on the neurocognitive processes of equal choices, while participants maintained a clear understanding of cue values for rational decisions between unequal options.

We first examined how reward certainty influences behavior, and whether a preference bias was present. We then fitted an accumulator model (Brown and Heathcote, 2008) to the behavioral performance across reward certainty levels. Posterior model parameters were used to infer whether the behavioral effects related to the decision process or visuomotor latencies unrelated to decision-making. EEG data were analyzed with time-resolved multivariate pattern classification for decoding spatiotemporal representations of reward certainty and preference. To establish a direct link between the decision process and its EEG signatures, we integrated behavioral and EEG data into a joint hierarchical Bayesian model and tested the hypothesis that electrophysiological activity reflects trial-by-trial changes in the speed of evidence accumulation for decisions.

We demonstrated that reward certainty and spontaneous preference independently shape RT and choices during equal choices. These behavioral effects extended further to other types of decisions, affected the decision process and evoked distinct electrophysiological patterns. Our findings highlighted that voluntary decisions are not random but subjected to exogenous and endogenous control.

## Materials and Methods

### Participants

Twenty-three healthy participants were recruited from Cardiff University School of Psychology participant panel (20 females; age range 19-32, mean age 22.7 years; 22 right-handed). All participants had normal or corrected-to-normal vision, and none reported a history of neurological or psychiatric illness. Written consent was obtained from all participants. The study was approved by the Cardiff University School of Psychology Research Ethics Committee.

### Apparatus

The experiment was conducted in a dedicated EEG testing room. A computer was used to control visual stimulus delivery and record behavioral responses. Visual stimuli were presented on a 24-inch LED monitor (ASUS VG248) with a resolution of 1920 by 1080 pixels and a refresh rate of 60 Hz, located approximately 100 cm in front of participants. Participants’ responses were collected from a response box (NATA technologies). The experiment was written in Matlab (Mathworks; RRID: SCR_001622) and used the Psychophysics Toolbox Version 3 extensions (Brainard, 1997; Kleiner et al., 2007, RRID: SCR_002881).

### Experimental design

All participants performed a decision-making task with probabilistic rewards during EEG recording (Fig. 1A). Before the decision task, the participants memorized 6 unambiguous cues in different shapes and their associated probabilities of receiving a reward (Fig. 1B and see *Procedures*). All the cues had the same color (RGB = 246, 242, 92) on a black background (100% contrast). Each cue was mapped onto one of the three reward certainty levels: high (a reward probability of 100%, i.e., always rewarded), medium (a reward probability of 80%) and low (a reward probability of 20%), and hence there were two different cues associated with each reward probability.

**Figure 1.**
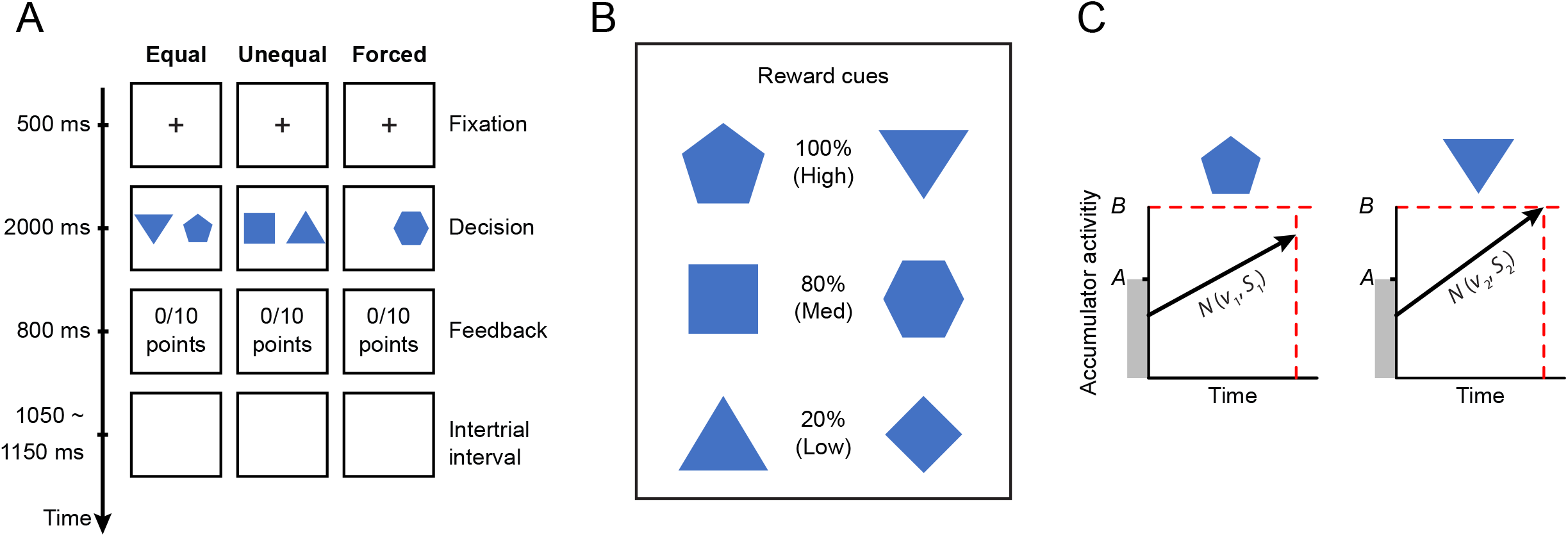
(**A**) Experimental paradigm of the probabilistic reward task. Participants were instructed to decide between two reward cues (equal and unequal trials) or respond to a single cue (forced trials). (**B**) A total of six reward cues were randomly assigned to three levels of reward certainty (100%, 80% or 20%). (**C**) Exemplar time course of the linear ballistic accumulator (LBA) model for equal choices. On each trial, the LBA assumes that evidence for two options are accumulated linearly and independently over time in two accumulators. The accumulation rate is sampled from a normal distribution with mean *v* and standard deviation *S*. The starting point of the accumulation process is sampled from a uniform distribution between 0 and *A*. The accumulation process terminates once the accumulated evidence first reaches a threshold *B*, and a corresponding decision is made by the winning accumulator.

The participants were instructed to maximize the total accumulated reward in the decision-making task. The task contained three conditions of trials: *equal*, *unequal* and *forced*. On an *equal* trial, two different cues with the same reward certainty appeared on the left and right sides of a central fixation point (e.g., 100% vs. 100%, 80% vs. 80% or 20% vs. 20%). On an *unequal* trial, two cues with different reward certainty levels appeared on both sides of the central fixation point (e.g., 100% vs. 20%, 100% vs. 80% or 80% vs. 20%). On a *forced* trial, one of the six cues appeared on either the left or right side of the fixation point. In equal and unequal trials, participants chose the left or right cue via button presses with the right-hand index and middle fingers. In forced trials, the participants responded to which side the single cue was presented (i.e., left or right). In all trials, the reward was operationalized as 10 virtual “game points” that did not have any tangible value. The probability of receiving the reward in a trial was either 100%, 80% or 20%, which was determined by the chosen cue. It is worth noting that, in equal trials, participants’ decisions did not actually affect the probability of receiving the reward because both options had equal reward certainty. In forced trials, if the participants chose the wrong side with no cue presented (0.1% across all forced trials), no reward was given. Feedback of rewarded (a “10 points” text message on the screen) or not rewarded (blank screen) was given after each trial. The total game points awarded were presented at the bottom of the screen throughout the experiment.

### Procedure

Each experimental session comprised 640 trials, which were divided into 4 blocks of 160 trials. Participants took short breaks between blocks and after every 40 trials within a block. The mappings between the six reward cues and three levels of reward certainty were randomized across participants. During breaks, the cues-reward mappings were explicitly presented on the screen (Fig. 1B), and the participants could take as much time as they needed to memorize them. After the first two blocks, all the cues were re-mapped to different reward probabilities. For example, if a cue was associated with 100% reward probability in the first and second blocks, the same cue would be associated with either 80% or 20% reward probability in the third and fourth blocks. The participants were encouraged to memorize the altered cue-probability associations prior to the third block. This re-mapping procedure reduced the potential bias associated with specific cue shapes.

Each block contained 64 equal trials (32 for 100% vs. 100%, 16 for 80% vs. 80% and 16 for 20% vs. 20%); 64 unequal trials (32 for 80% vs. 20%, 16 for 100% vs. 80% and 16 for 100% vs. 20%) and 32 forced trials (16 for *100%*, 8 for *80%* and 8 for *20%*) at a randomized order. This design ensured the same number of trials with and without cues with the highest reward certainty (100%). Because there were two cues at each level of reward probability, different cue combinations and their positions on the screen can give the same pair of reward probability (e.g., there are 8 possible combinations for 80% vs. 20% unequal trials). These different combinations were counterbalanced across trials.

Each trial began with the presentation of a fixation point at the center of the screen, which was presented for 500 ms. After the fixation period, in the equal and unequal trials, two reward cues appeared on the left and right sides of the screen with a horizontal distance of 4.34° from the fixation point. Both cues were vertically centered. In forced trials, only one reward cue appeared on one side of the screen, and the side of cue appearance was randomized and counterbalanced across trials. Cues were presented for a maximum of 2000 ms, during which the participants were instructed to make a left or right button press. The cues disappeared as soon as a response was made, or the maximum duration was reached. The reaction time (RT) on each trial was measured from the cue onset until participants made a response. Feedback (rewarded or not rewarded) was given 200 ms after the reward cue offset and lasted 800 ms, followed by a random intertrial interval uniformly distributed between 1050 and 1150 ms. As in our previous study (Zhang and Rowe, 2014), if the participant failed to respond within 2000 ms or responded within 100 ms, no reward was given and a warning message “*Too slow*” or “*Too fast*” was presented for 1500 ms.

### Behavioral analysis

We excluded trials with RT faster than 200 ms (fast guesses). For each participant, trials with RTs longer than 2.5 standard deviations from the mean RT were also excluded from subsequent analysis. The discarded trials accounted for 1.5% of all trials.

We first analyzed the proportion of choices in equal trials to establish the existence of a preference bias. In the equal condition, by definition, there was no “correct” or “incorrect” response, because two cues in an equal trial had the same reward probability. For each pair of cues with the same reward probability, we defined the *preferred* cue as the one chosen more frequently than the other (*non-preferred*) one in equal trials. The categorization of preferred and non-preferred cues was estimated separately between the first two and the last two blocks, because of the cue-probability remapping after the first two blocks. At each level of reward certainty, a *preference bias* was then quantified as the proportion of trials where the preferred cue was chosen. The preference bias had a lower bound of 50%, at which there was no choice bias between the two cues with equal reward certainty.

In the unequal condition, we defined the decision accuracy as the proportion of choosing the cue with higher reward probability, separately for each combination of reward probabilities (100% vs. 80%, 100% vs. 20% and 80% vs. 20%). Two-tailed one-sample t-tests compared the decision accuracy in the unequal condition against a chance level of 50%, which would indicate non-rational decisions (i.e., both high and low reward cues were chosen in 50% of trials).

To determine how reward certainty, preferences and other experimental factors influence RT, we analyzed single-trial RT data with linear mixed-effects models (LMMs) using the lme4 package (Bates et al., 2015b) in R (RRID: SCR_001905). The LMM is a hierarchical regression method that distinguishes between fixed and random effects (Gueorguieva and Krystal, 2004). LMMs take into account all single-trial data without averaging across trials and offer better control of type 1 and type 2 errors than ANOVA (Baayen et al., 2008). Therefore, statistical inferences from LMMs are robust to experimental designs with unbalanced trials across conditions (Bagiella et al., 2000), which is an important feature suitable for the current study.

We designed two LMMs with different dependent variables and factors (Table 1). Model 1 analyzed the RTs from equal and forced trials, including choice type (equal or forced) and reward certainty (high, medium or low) as factors. For the unequal condition, because each trial had two cues with different levels of reward certainty that cannot be directly compared with equal or forced trials, the RTs of unequal trials were analyzed separately in Model 2. The sum and the absolute difference of the two reward probabilities in each unequal trial were included as factors of RT, as they have been shown to affect choice behavior (Thaler, 1981; Teodorescu et al., 2016; Ballard et al., 2017).

**Table 1.**
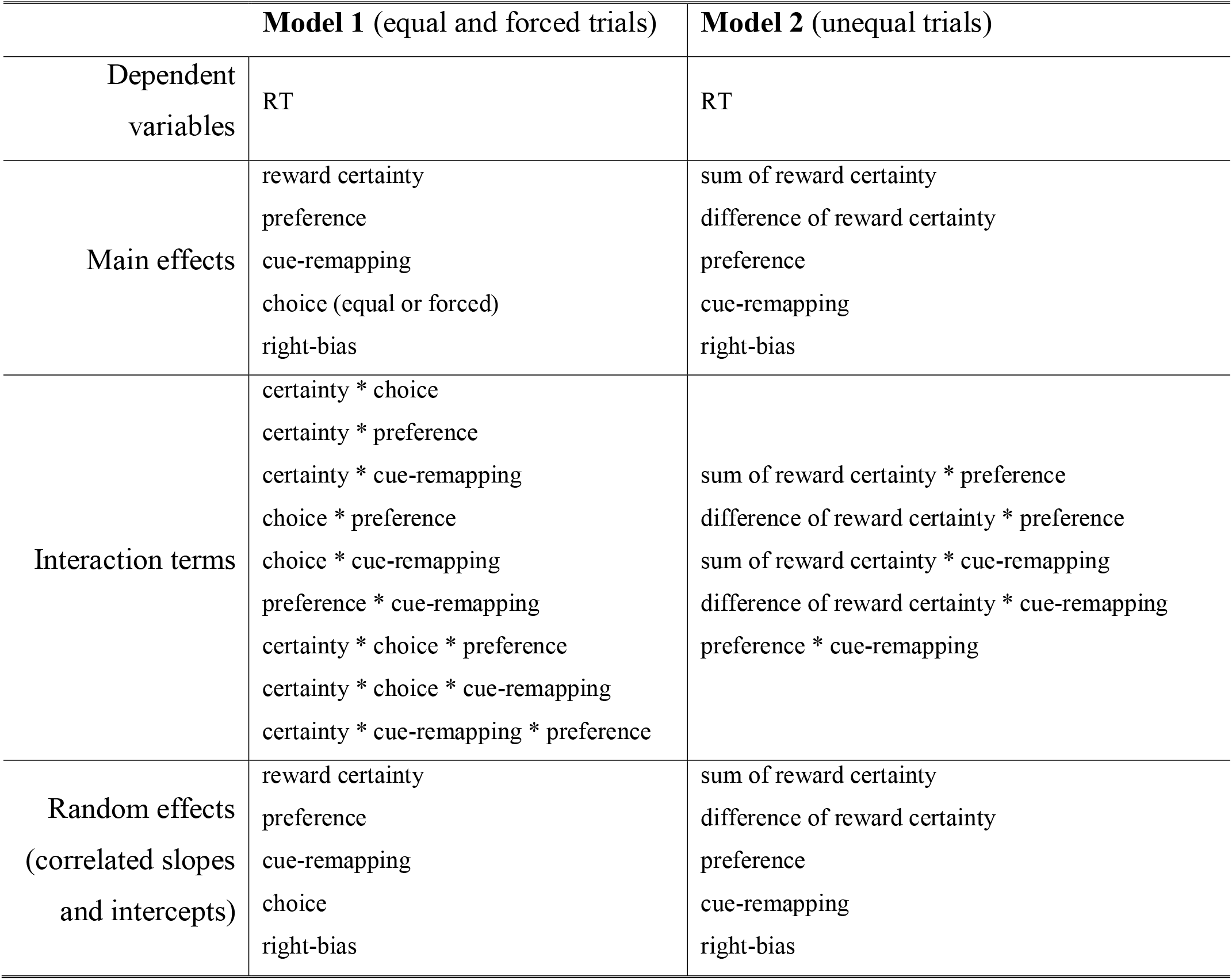
The linear mixed-effects models of RT. Model 1 analyzed single-trial RT in equal and forced trials. Model 2 analyzed single-trial RT in unequal trials. In both models, *preference* was a predictor indicating whether the preferred cue was selected in each trial. *Cue-remapping* was a predictor indicating whether each trial was before or after cure-reward re-mapping in the second half of each session. *Right-bias* indicated whether the cue on the right size of the screen was chosen in each trial, modeling a possible response bias.

|  | **Model 1** (equal and forced trials) | **Model 2** (unequal trials) |
| --- | --- | --- |
| Dependent variables | RT | RT |
| Main effects | reward certainty<br>preference<br>cue-remapping<br>choice (equal or forced)<br>right-bias | sum of reward certainty<br>difference of reward certainty<br>preference<br>cue-remapping<br>right-bias |
| Interaction terms | certainty * choice<br>certainty * preference<br>certainty * cue-remapping<br>choice * preference<br>choice * cue-remapping<br>preference * cue-remapping<br>certainty * choice * preference<br>certainty * choice * cue-remapping<br>certainty * cue-remapping * preference | sum of reward certainty * preference<br>difference of reward certainty * preference<br>sum of reward certainty * cue-remapping<br>difference of reward certainty * cue-remapping<br>preference * cue-remapping |
| Random effects<br>(correlated slopes<br>and intercepts) | reward certainty<br>preference<br>cue-remapping<br>choice<br>right-bias | sum of reward certainty<br>difference of reward certainty<br>preference<br>cue-remapping<br>right-bias |

In all the LMMs, fixed effects structures included hypothesis-driven, design-relevant factors and their interactions, and individual participants were included as the source of random variance (random effect). We used a standard data-driven approach to identify the random effects structure justified by the experimental design, which resulted in good generalization performance (Barr et al., 2013; bust see Bates et al., 2015 for alternative methods). This approach starts with the maximal random effects structure (i.e., including all random slopes, intercepts and interactions) and systematically simplified it until the LMM reaches convergence. Table 1 lists the simplified random effects structures. The correlation structures of each fitted LMM was assessed to avoid overfitting (Matuschek et al., 2017).

### A cognitive model of voluntary decision-making

We further analyzed the behavioral data using a cognitive model, the Linear Ballistic Accumulator (LBA) model (Brown and Heathcote, 2008). The LBA model is a simplified implementation of a large family of sequential sampling models of decision-making (Ratcliff and Smith, 2004; Bogacz et al., 2006; Gold and Shadlen, 2007; Zhang, 2012). It has been used to examine cognitive processes during perceptual decision (Ho et al., 2009; Forstmann et al., 2010a) voluntary choice (Zhang et al., 2012) and action initiation (Karahan et al., 2019).

Our model-based analysis served three purposes in the current study. First, we fitted a family of LBA models with various model complexity to behavioral data of individual participants in equal trials. By identifying the best-fitted model, we inferred how reward certainty and preference modulated subcomponents of the evidence accumulation process during decision-making. Second, we simulated the best fitted LBA model and examined whether model simulations were consistent with the experimental data in forced and unequal conditions. This is a stringent test of model generalizability because the experimental data in forced and unequal trials were unseen by the model fitting procedure. Third, we linked the cognitive processes identified by the LBA model to brain activities. This was achieved by incorporating a trial-by-trial measure of EEG activity regressors into the best-fitted model (Cavanagh et al., 2011; Nunez et al., 2017, 2018, and see *EEG-informed cognitive modelling*).

In equal trials, the participants made binary choices between two cues associated with the same reward certainty. The LBA model assumed that the decision of *when* and *which* to choose is governed by a “horse race” competition between two accumulators *i* ∈{1, 2} that accumulate evidence over time supporting the two choice options (Fig. 1C). One accumulator is in favor of the preferred cue, and the other accumulator is in favor of the non-preferred cue. The activations of the accumulators represent the accumulated evidence. At the beginning of each trial, the initial activations of the two accumulators are independently drawn from a uniform distribution between 0 and *A*. The activation of each accumulator then increases linearly over time, and the speed of accumulation (i.e., accumulation rate) varies as a Gaussian random variable with mean *v_i_* and standard deviation *S_i_* across trials. The accumulation process terminates when the activation of any accumulator reaches a response threshold *B* (*B* > *A*) and the choice corresponding to the winning accumulator is selected. The model prediction of RT is the sum of the duration of the accumulation process and a constant non-decision time *T*_er_, with the latter accounts for the latency associated with other processes including stimulus encoding and action execution (Brown and Heathcote, 2008; Nunez et al., 2018; Karahan et al., 2019).

### Model parameter estimation and model selection

The LBA model has five key parameters: the mean *v* and standard deviation *S* of the accumulation rate across trials, the decision threshold *B*, the standing point variability *A* and the non-decision time *T*_er_. To accommodate the empirical data, one or more model parameters need to vary between conditions. We evaluated a total of 21 variants of the LBA model with different parameter constraints (Fig. 3A). First, the accumulation process may differ between the preferred and non-preferred options, leading to *v* or *S* to vary between accumulators (preferred, non-preferred). Second, reward certainty could modulate the accumulation process or visuomotor latencies unrelated to decisions, leading to *v*, *S* or *T*_er_ to vary between three levels of reward certainty (100%, 80% and 20%). Third, the decision threshold *B* was fixed between conditions, because the trial order was randomized, and we do not expect the participants to systematically vary their decision threshold before knowing the cues to be presented (Ratcliff and Smith, 2004). During model-fitting, the decision threshold was fixed at 3 as the scaling parameter (Brown and Heathcote, 2008), and all the other parameters allowed to vary between participants. Finally, because the participants showed behavioral differences between reward certainty levels and between preferred/non-preferred choices, we only estimated realistic models: those with at least one parameter varied between reward certainty levels (*v*, *S* or *T*_er_) and at least one parameter varied between accumulators (*v* or *S*).

We used a hierarchical Bayesian model estimation procedure to fit each LBA model variant to individual participant’s choices (the proportion of preferred and non-preferred choices) and RT distributions in equal trials. The hierarchical model assumes that model parameters at the individual-participant level are random samples drawn from group-level parameter distributions. Given the observed data, Bayesian model estimation uses Markov chain Monte Carlo (MCMC) methods to simultaneously estimate posterior parameter distributions at both the group level and the individual-participant level. The hierarchical Bayesian approach has been shown to be more robust in recovering model parameters than conventional maximum likelihood estimation (Jahfari et al., 2013; Zhang et al., 2016).

For group-level parameters (*v*, *S*, *A* and *T*_er_), similar to previous studies (Annis et al., 2017), we used weakly informed priors for their means *E*(.) and standard deviations *std*(.):

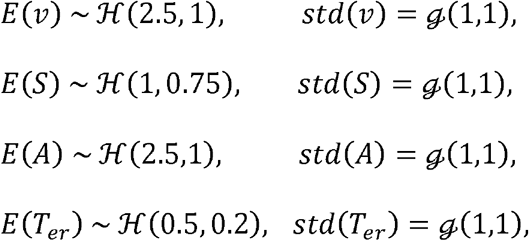

where 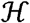 represents a positive normal distribution (i.e., truncated at 0) with parameterized mean and standard deviation, and 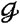, represents a Gamma distribution with parameterized mean and standard deviation.

We used the *hBayesDM* package (Ahn et al., 2017) in *R* for the hierarchal implementation of the LBA model. For each of the 21 model variants, we generated four independent chains of 7,500 samples from the joint posterior distribution of the model parameters using Hamiltonian Monte Carlo (HMC) sampling in Stan (Carpenter et al., 2017). HMC is an efficient method suitable for exploring high-dimensional joint probability distributions (Betancourt, 2017). The initial 2,500 samples were discarded as burn-in. To assess the convergence of the Markov chains, we calculated Gelman-Rubin convergence diagnostic 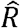 of each model (Gelman and Rubin, 1992) and used 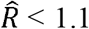 as a stringent criterion of convergence (Annis et al., 2017). We compared the fitted LBA model variants using Bayesian leave-one-out information criterion (LOOIC). LOOIC evaluates the model fit while considering model complexity, with lower values of LOOIC indicating better out-of-sample model prediction performance (Vehtari et al., 2017).

### Model Inferences

We used Bayesian inference to analyze the posterior distributions of group-level model parameters (Berger and Bayarri, 2004). To evaluate if a parameter varies substantially between two conditions, we calculated the proportion of posterior samples in which the parameter value for one condition was greater than the other. To test if a parameter differs from a threshold value, we calculated the proportion of the posteriors greater or smaller than the threshold. To avoid confusion, we used *p* to refer to classical frequentist *p*-values, and *P_p|_*_D_ to refer to Bayesian inference results based on the proportion of posteriors supporting the testing hypothesis, given the observed data.

### Model simulations

To evaluate the model fit to the empirical data of equal choices, we calculated the posterior prediction of the best fitted LBA model by averaging 100 iterations of model simulation using posterior parameter estimates. Averaging across multiple iterations reduces potential biases when sampling from posterior parameter estimates. Each of the 100 iterations generated simulated behavioral responses (i.e., RTs and choices) of individual participants, with the same number of trials per condition as in the actual experiment.

We further used the best fitted LBA model to simulate unequal and forced choice responses. This allowed us to evaluate whether the model fitted to the equal choice data could characterize behavioral patterns in other types of choices. For unequal choices, two accumulators representing two cues with different reward certainty levels compete to reach the decision threshold, with their parameters set to the posterior estimates from the fitted LBA model. For forced choices, a single accumulator was set to reach to the decision threshold. Similar to the simulation of equal choices, for each participant, we averaged the predicted behavioral responses of unequal and forced choices from 100 iterations of simulation. Each iteration contained the same number of trials as in the experiment.

### EEG data acquisition_and processing

EEG data were collected using a 32-channel Biosemi ActiveTwo device (BioSemi, Amsterdam). Due to technical issues, EEG data collection was not successful in two participants, and therefore all EEG data analyses were performed on the remaining 21 participants. EEG electrodes were positioned at standard scalp locations from the International 10-20 system. Vertical and horizontal eye movements were recorded using bipolar electrooculogram (EOG) electrodes above and below the left eye as well as from the outer canthi. Additional electrodes were placed on the mastoid processes. EEG recordings (range DC-419 Hz; sampling rate 2048 Hz) were referenced to linked electrodes located midway between POz and PO3/PO4 respectively and re-referenced off-line to linked mastoids.

EEG data were pre-processed using EEGLab toolbox 13.4.4b (Delorme and Makeig, 2004, RRID: SCR_007292) in Matlab. The raw EEG data were high-pass filtered at 0.1 Hz, low-pass filtered at 100 Hz using Butterworth filters and downsampled to 250 Hz. An additional 50 Hz notch filter was used to remove mains interference. We applied Independent Component Analysis (ICA) to decompose continuous EEG data into 50 spatial components, using *runica* function from the EEGLab toolbox. Independent components reflecting eye movement artifacts were identified by the linear correlation coefficients between the time courses of independent components and vertical and horizontal EOG recordings. Additional noise components were identified by visual inspection of the components’ activities and scalp topographies. Artefactual components were discarded, and the remaining components were projected back to the data space.

After artifact rejection using ICA, the EEG data were low-pass filtered at 40 Hz and epoched from −400 ms to 1000 ms, time-locked to the onset of the stimulus (i.e., reward cues) in each trial. Every epoch was baseline corrected by subtracting the mean signal from −100 ms to 0 ms relative to the onset of reward cues.

### Event_related potential (ERP) analysis

We examined univariate differences in evoked responses between conditions in single EEG electrodes. For each participant, trial-averaged ERPs were calculated from epochs of equal or forced choices. For both equal and forced conditions, we tested for differences in ERPs between three levels of reward certainty using a one-way repeated-measures ANOVA. Furthermore, we tested for differences in ERPs between preferred and non-preferred choices in equal trials, using a paired t-test. We performed statistical tests in all electrodes and all time points. Cluster-based permutation tests (2000 iterations with maximum statistics) were used to correct for multiple comparisons across electrodes and time points (Maris and Oostenveld, 2007).

### Multivariate pattern analysis

We used time-resolved MVPA on pre-processed, stimulus-locked EEG data to assess reward-specific and preference-specific information throughout the time course of a trial. In contrast to univariate ERP analysis, MVPA combines information represented across multiple electrodes, which has been shown to be sensitive in decoding information representation from multi-channel human electrophysiological data (Cichy et al., 2014; Dima et al., 2018).

We conducted three MVPA analysis to identify the latency and spatial distribution of the EEG multivariate information. The first was to decode reward certainty levels in equal choices (e.g., equal trials with two 100% reward cues versus equal trials with two 80% cues). The second was to decode preferred versus non-preferred choices in equal trials. The third was to decode between equal and forced choices with the same reward certainty (e.g., equal trials with two 100% cues versus forced trials with a 100% cue).

Each analysis was formed as one or multiple binary classification problems, and the data feature for classification included EEG recordings from all 32 electrodes. In each analysis, at each sampled time point (−400 ms to 1000 ms) and for each participant, we trained linear support vector machines (SVM) (Garrett et al., 2003) using the 32-channel EEG data and calculated the mean classification accuracy following a stratified ten-fold cross-validation procedure. In all MVPA, we included the EEG data from 400 ms before cue onset as a sanity check, because one would not expect significant classification before the onset of reward cues.

In each cross-validation, 90% of the data was used as a training set, and the remaining 10% of the data was used as a test set. In some analysis (e.g., equal trials with 100% cues versus equal trials with 80% cues), the number of samples belonging to the two classes was unbalanced in the training set. We used a data-driven over-sampling approach to generate synthetic instances for the minor class until the two classes had balanced samples (Zhang and Wang, 2011). The synthetic instances were generated from Gaussian distributions with the same mean and variance as in the original minority class data. Training set data were standardized with z-score normalization to have a standard normal distribution for each feature. The normalization parameters estimated from the training set was then applied separately to the test set to avoid overfitting. To reduce data dimensionality, we performed principal component analysis to the training set data and selected the number of components that explained over 99% of the variance in the training set. The test set data were projected to the same space with reduced dimensions by applying the eigenvectors of the chosen principal components. We then trained SVM to distinguish between the two classes (i.e., conditions) and evaluate the classification accuracy using the test set data. The procedure was repeated ten times with different training and test sets, and the classification accuracies were averaged from the ten-fold cross-validation. We used the SVM implementation in MATLAB Machine Learning and Statistics Toolbox. The trade-off between errors of the SVM on training data and margin maximization was set to 1.

To estimate the significance of the classification performance, we used two-tailed one-sample t-test to compare classification accuracies across participants against the 50% chance level. To account for the number of statistical tests at multiple time points, we used cluster-based permutation (Maris and Oostenveld, 2007) to control the family-wise error rate at the cluster level from 2000 permutations.

### Spatial patterns of classification weights

To evaluate the relative importance of each feature (i.e., EEG electrode) to the classification performance, we calculated the weight vector of SVMs. For each classification problem, we retrained the SVM at each time point with all the data included in the training set and obtained the SVM weight vector. The weight vectors were then transformed into interpretable spatial patterns by multiplying the data covariance matrix (Haufe et al., 2014). The group spatial patterns were calculated by averaging across participants and from all time points which had significant classification accuracy.

### Estimation of single-trial ERP components

We estimated two ERP components from single-trial EEG data in equal trials: N100 and P300, which were subsequently used to inform cognitive modelling. The visual N100 is related to visual processing (Mangun and Hillyard, 1991) and the P300 is related to evidence accumulation during decision making (Kelly and O’Connell, 2013; Twomey et al., 2015).

To improve the signal-to-noise ratio of single-trial ERP estimates, we used a procedure similar to previous studies (Kayser and Tenke, 2003; Parra et al., 2005; Nunez et al., 2018). For each participant, we first performed singular value decomposition (SVD) to the grand averaged ERP data across all equal trials. The first SVD component is the one that explained the most variance in the averaged ERP waveforms, and its weight vector provides an optimal spatial filter to detect the ERP waveforms. Next, we applied the optimal spatial filter as a channel weighting function to single-trial EEG data.

The single-trial EEG data filtered with the SVD-based weighting function was then used to identify the peak-latency and peak-amplitude of the N100 and P300 components. For N100, we searched for the peak negative amplitude in a window centered at the group-level N100 latency (112 ms) and started at 60 ms. The lower bound of the search window was determined by the evidence that the visual onset latency is ∼60 ms in V1 (Schmolesky et al., 2017). For P300, we searched for a peak positive amplitude in a window centered at the group-level P300 latency (324 ms). For both N100 and P300, the search window had a length of 104 ms, similar to a previous study (Nunez et al., 2018).

### EEG-informed cognitive modeling

From the behavioral responses of each participant and each condition, we used the LBA model to estimate a fixed mean accumulation rate across trials (see *Model parameter estimation and model selection*). Recent studies showed that the variability of the P300 component closely relates to the rate of evidence accumulation during decision making (Twomey et al., 2015). We therefore extended the best fitted LBA model with EEG-informed, single-trial regressors, which estimated the effect of trial-by-trial variability in EEG activity on the mean accumulation rate (Hawkins et al., 2015; Nunez et al., 2017).

The main regressor of interest was the slope of change between the P300 and N100 components, which were estimated as the ratio of the P300-N100 peak-amplitude difference and the P300-N100 peak-latency difference in each equal trial. We also tested four additional regressors from individual ERP components: P300 amplitude, P300 latency, N100 amplitude and N100 latency. All the EEG regressors were obtained from the estimations of single-trial ERP components in equal choice trials. To obtain a meaningful intercept, the regressors were mean-centered and rescaled to have a unit standard deviation.

Each EEG regressor was tested in a linear regression model, using the same Bayesian hierarchical model estimation procedure as in the behavioral modelling analyses. For each regression model, we assumed that the mean accumulation rates of both accumulators *v*_1_(*t*) and *v*_2_(*t*) (i.e., the one in favor of the preferred option and the other one in favor of the non-preferred option) are influenced by the EEG regressor of interest on a trial-by-trial basis:

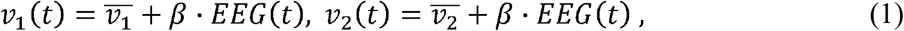

while *t*=1, 2, 3, … represents the equal choice trials, and 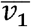 and 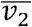 are the intercepts. The regression coefficient *β* represents the effect of EEG regressor on the mean accumulation rates.

### Software and data accessibility

The scripts for behavioral modelling and EEG analysis are open-source and freely available online (https://github.com/ccbrain/voluntary-decision-eeg). We have also made the behavioral and EEG data open access (https://doi.org/10.6084/m9.figshare.9989552.v1).

## Results

We examined the effects of reward certainty and spontaneous preference on behavior and EEG activity during voluntary decisions. In a probabilistic reward task, participants chose between two options with the same reward certainty (equal trials) at high (100%), medium (80%) or low (20%) levels. In additional control conditions, the participants made binary choices between options with different levels of reward certainty (unequal trials) or responded to the location of a single reward cue (forced trials). Below, we first reported behavioral results. We then fitted LBA models (Brown and Heathcote, 2008) to the choices and RT distributions of equal trials and make inferences about the model parameters that quantify an evidence accumulation process during voluntary decisions. Next, we performed univariate and multivariate analyses of EEG data to identify spatiotemporal representations of reward certainty and preference information as well as their time courses. We then extended the best-fitted LBA model with single-trial measures of EEG activity to test whether trial-to-trial variations in EEG data relates to the rate of evidence accumulation across trials.

### Behavioral results: choices

In equal choice trials, we calculated the preference bias as the proportion of choosing the cue with higher frequency in equal trials (i.e., the preferred cue). At each level of reward certainty, there was a strong preference bias (>50%) for choosing one reward cue over the other (Fig. 2A; high: 95% CI [0.682, 0.765]; medium: 95% CI [0.679, 0.759]; low: 95% CI [0.669, 0.745]). A repeated-measures ANOVA showed no significant difference in preference bias between reward certainty levels (*F*(2, 44) = 0.2, *p* = 0.81). Therefore, although the two options had the same reward certainty, participants did not make their choices randomly. Because the cue-probability mapping was randomized across participants and re-mapped within each session, the observed preference bias could not be readily explained by a group-level preference towards any specific cue, but rather a spontaneous bias at the individual level.

**Figure 2.**
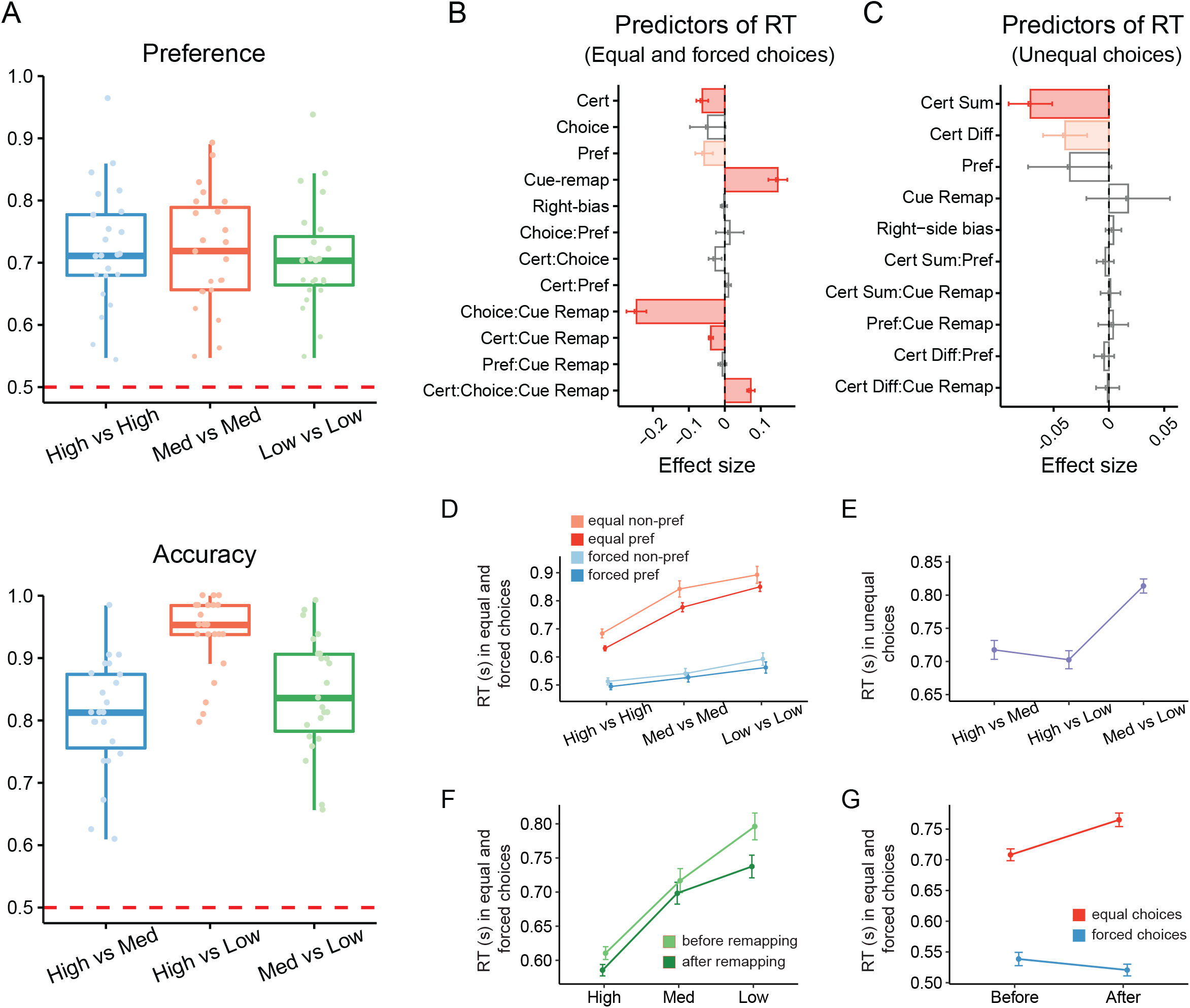
Behavioral results. (**A**) Preference bias across reward certainty levels in equal trials (top) and decision accuracy across reward certainty levels in unequal trials (bottom). (**B**) Linear mixed-effects model results for Model 1 in Table 1. Dark red bars represent significant effects with *p* < 0.001. Light red bars represent significant effects with *p* < 0.05. Grey bars represent non-significant factors and interactions. Error bars represent standard errors across participants. (**C**) Linear mixed-effects model results for Model 2 in Table 1. Significant effects and interactions in RT from Model 1 (Table 1) were presented separately for: reward certainty and preference in equal and forced trials (**D**), before and after cue-remapping at different reward certainty levels (**F**), before and after cue-remapping in equal and forced trials (**G**). Significant main effects in Model 2 were presented in panel **E**. In panels D-G, Error bars represent standard errors across participants.

In unequal choice trials, as expected, the cues with higher reward certainty levels were chosen more often, as evidenced by the above-chance decision accuracies in all conditions (Fig. 2A; high vs. medium: *t*(22) = 16.08, 95% CI [0.774, 1], *p* < 0.001; high vs. low: *t*(22) = 23.31, 95% CI [0.862, 1], *p* < 0.001; medium vs. low: *t*(22) = 20.97, 95% CI [0.834, 1], *p* < 0.001; one-sample t-test against the 0.5 chance level). A repeated-measures ANOVA showed significant differences in decision accuracy between reward certainty levels (*F*(2, 44) = 28.17, *p* < 0.001).

Post-hoc pairwise comparison with Tukey’s correction indicated that accuracy in the high vs. low certainty condition (93.8%) was significantly higher than in the high vs. medium (84.3%; *t*(44) = 5.267, *p* < 0.001) and the medium vs. low (80.7%; *t*(44) = 7.265, *p*<0.001) conditions. These results suggested that the participants remembered the cue-reward mapping for rational choice behavior.

### Behavioral results: response time

We used a linear mixed-effects model (LMM) to quantify the influence of experimental factors on RTs in equal and forced choices (Fig. 2B, Model 1 in Table 1). The fixed effects included reward certainty, choice type (equal vs. forced), preference (choosing the preferred vs. the non-preferred option), cue re-mapping (before vs. after the cue-reward remapping halfway through each session) and their meaningful interactions (Figs. 2D-2F). The LMM included individual participants as random intercepts and data-driven random effects structures (Barr et al., 2013). The participants were faster in responding with the preferred than the non-preferred option (Fig. 2D, *β* = −0.063, 95% CI [−0.027, −0.991], *p* < 0.001) and RTs decreased as the reward certainty increased (*β* = −0.101, 95% CI [−0.067, −0.135], *p* < 0.001). The RT in equal choice trials were longer than that in forced trials (*β* = −0.292, 95% CI [−0.201, −0.384], *p* < 0.001). The effect of reward certainty on RT was stronger in equal choices than that in forced choices, supported by a significant interaction between the two main effects (*β* = 0.045, 95% CI [0.025, 0.066], *p* < 0.001).

Participants had slower responses after memorizing a new set of cue-reward associations, indicated by a significant main effect in RT before and after cue re-mapping (*β* = 0.149, 95% CI [0.096, 0.201], *p*<0.001). The significant interaction between cue re-mapping and reward certainty suggested that the increase in RT was more pronounced in trials with lower reward certainty (Fig. 2F, **β** = −0.039, 95% CI [−0.051, −0.026], *p* < 0.001). The interaction between cue re-mapping and choice type (Fig. 2G, *β* = −0.247, 95% CI [−0.192, −0.302], *p* < 0.001) further indicated that this pattern was mainly associated with equal choices than forced responses. Because evaluating the reward certainty of a cue was likely associated with additional cognitive load after cue re-mapping, the observed RT difference before and after cue re-mapping implies that, in equal choices, the participants evaluated both cues throughout the experimental session.

In a second LMM, we analyzed RTs in unequal trials (Model 2 in Table 1), including the sum and difference of the reward certainty of two cues in each trial as fixed effects. The sum of the two reward probabilities in unequal trials was negatively associated with RT (Fig. 2E, *β* = −0.071, 95% CI [−0.032, −0.110], *p* < 0.001), consistent with previous studies that the total reward magnitude influences decision-making (Pirrone et al., 2014; Teodorescu et al., 2016).

### Cognitive modelling of behavioral data

We showed that the behavioral performance in equal choices was modulated by both reward certainty and spontaneous preference. Similar to previous studies (Forstmann et al., 2010b; Mulder et al., 2010, 2014; Zhang et al., 2012), we conceptualized that a decision between two equal options is described by an LBA model. The model assumes that the momentary evidence for choosing the preferred and non-preferred options accumulate independently over time, until a decision threshold is reached (Fig. 1C).

To identify the cognitive processes that led to the observed behavioral differences, we compared 21 variants of the LBA model. The model variants differed systematically in their constraints on whether the rate of evidence accumulation rate and the non-decision time could change between reward certainty levels or between preferred/non-preferred options. For each model variant, we used hierarchal Bayesian modelling with MCMC parameter estimation routine to estimate the posterior distributions of the model parameters, given the observed choice and RT distribution from individual participants (see *Model parameter estimation and model selection*). To identify the model with the best fit, we calculated the Bayesian leave-one-out information criterion (LOOIC) score of each model (Vehtari et al., 2017).

Posterior parameter estimates in all the 21 model variants were converged after sufficient samples (Gelman-Rubin convergence diagnostic 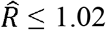 for all parameters in all models). The LOOIC scores suggested that the models with the mean accumulation rate varying between reward certainty levels and between preference levels fitted the data better than others model variants. The best-fitting model (i.e., the one with the lowest LOOIC score, Fig. 3A) had a fixed non-decision time per participant with the standard deviation of the accumulation rate varying between reward certainty levels and preferred/non-preferred options. To evaluate the model fit, we simulated the best fitting model and compared its posterior predictions with the observed equal choice data (Fig. 3B). There was a good agreement between the observed data and the model simulations across reward certainty levels and choice preferences.

**Figure 3.**
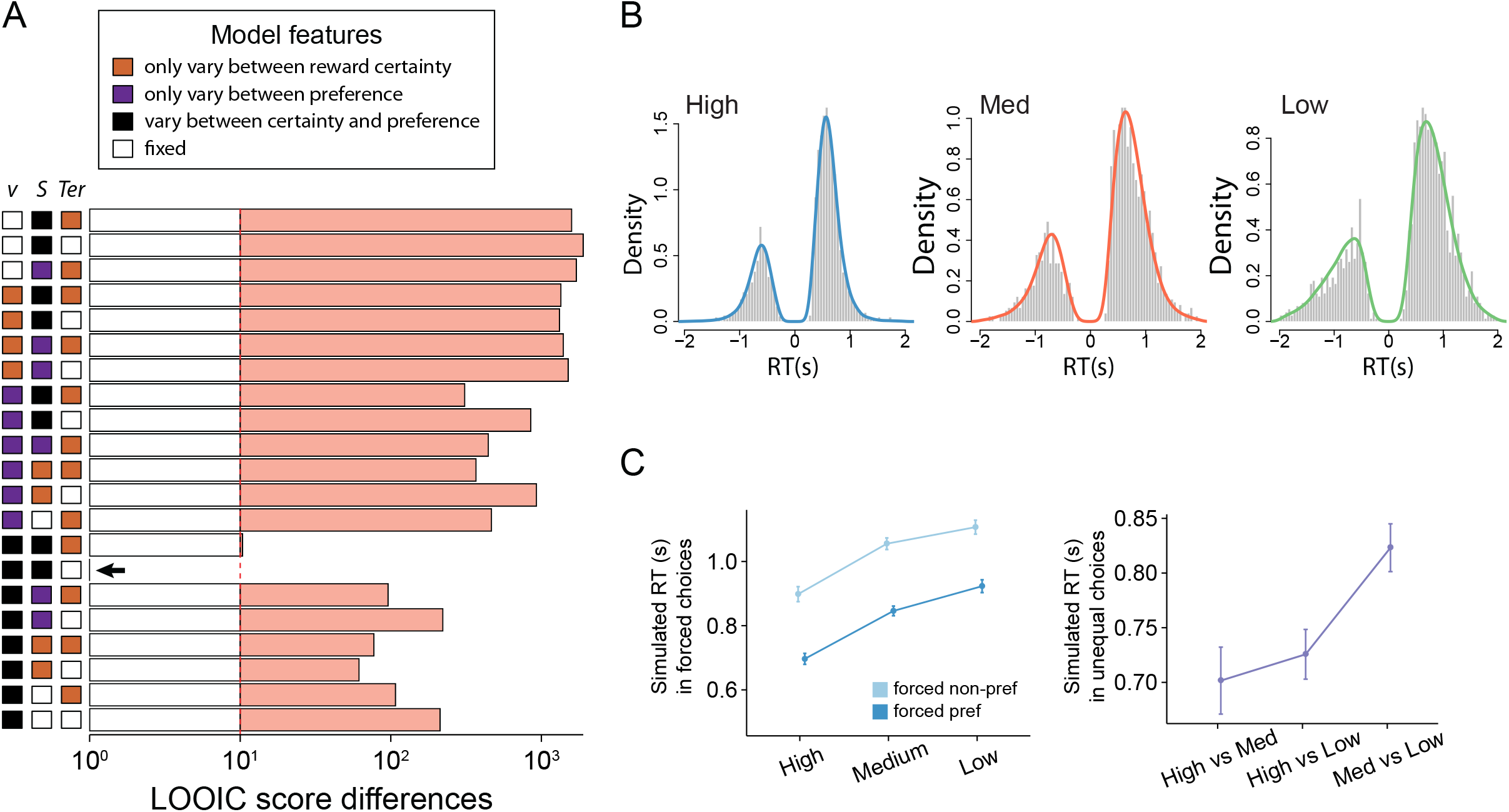
Model comparisons, model fits and model simulations. (**A**) LOOIC scores of 21 LBA model variants. The LOOIC score differences between all models and the best model are plotted against corresponding model structures, which were illustrated on the left of the panel. The model structure specified how the mean accumulation rate *v*, the standard deviation *S* of the accumulation rate and the non-decision time *T_er_* could vary between conditions. A black filled square indicated that the corresponding parameter could vary between reward certainty levels and preferred/non-preferred options. An orange or purple filled square indicated that the corresponding parameter could only vary between reward certainty levels or preferred/non-preferred options, respectively. Unfilled squares indicated that the parameter remained fixed between conditions. The best model was shown with a LOOIC score difference of zero (indicated by the arrow). (**B**) Simulations of RTs in equal choices, generated from the posterior distribution of the best fitted model for high (left), medium (middle) and low (right) reward certainty levels. Histograms represent experimental data and density distributions represent model simulation from 100 iterations. Negative values represent RTs for non-preferred choices. (**C**) Simulation of RTs in forced (left) and unequal (right) choices from 100 iterations. Error bars represent standard errors across participants.

For the best fitting model (Fig. 4A), we compared the posterior estimates of the group-level parameters between conditions (Fig. 4B and Fig. 4C). There was strong evidence for a larger reward certainty being associated with higher mean (*v*, high > low: *P_p|_*_D_ = 1; high > medium: *P_p|_*_D_ = 0.999; medium > low: *P_p|_*_D_ = 0.839) and standard deviation (*S*, high > low: *P_p|_*_D_ = 1; high > medium: *P_p|_*_D_ = 0.954; medium > low: *P_p|_*_D_ = 0.877) of the accumulation rate. There was also strong evidence for a higher mean accumulation rate for the preferred than the non-preferred options (*P_p|_*_D_ = 0.999), and no evidence for a difference in the standard deviation of the accumulation rate between preference levels (*P_p|_*_D_ = 0.532). These results supported that preferred cues and the ones with higher reward certainty were recalled and processed faster than non-preferred cues. The cues with higher reward certainty were also associated with more variable accumulation rate. Model comparisons further suggested that the latencies of early visual encoding and motor execution were not influenced by reward certainty nor preference, because the models with varying non-decision time had inferior fits.

**Figure 4.**
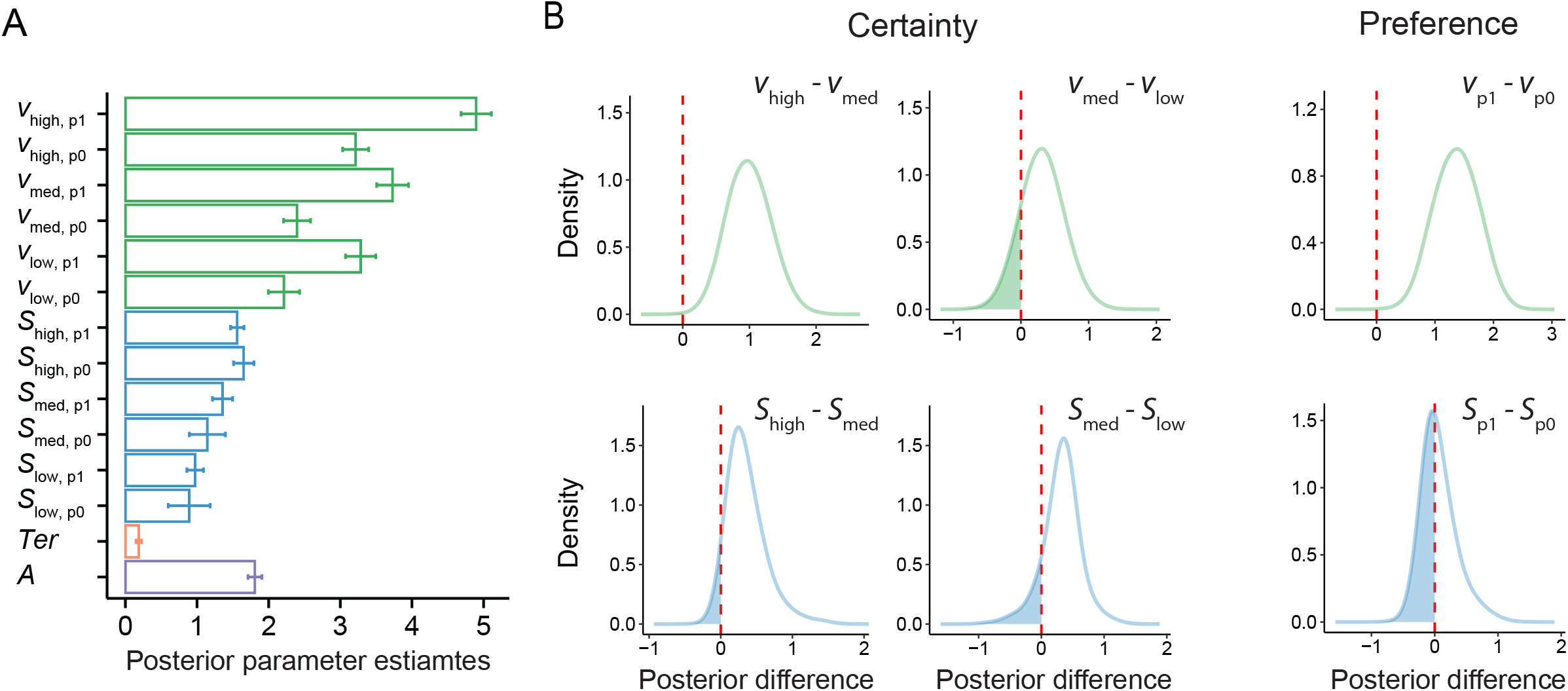
Posterior model parameters and inferences. (**A**) Group-level LBA model parameters of the best fitting model: means of accumulation rates (*v*, green), standard deviations of accumulation rates (*S*, blue), non-decision time (*T_eer_*; orange) and starting point (*A*, purple). Error bars represent standard deviations of posterior distributions of parameter values. The means and standard deviations of accumulation rates were shown separately for each reward certainty level (high, medium and low) and accumulator (p1, preferred option; p0, non-preferred option). (**B**) Differences of posterior parameter estimates across certainty levels (left and middle columns) and preference levels (right column). The proportion of posterior difference distributions above zero suggested higher parameter values for higher certainty level or more preferred options.

Next, we evaluated whether the best fitting model could reproduce qualitative RT patterns in the forced and unequal choices (Fig. 3C), which were unseen by the parameter estimation procedure (see *Model simulation*). For unequal choices, the simulated RT showed similar patterns to the observed data, in which choosing between medium and low certainty cues led to the longest RT. For forced choices, similar to the observed data, higher reward certainty and preferred cues were associated with faster RT in simulation. However, simulated RT in forced choices was longer than the experimental data, suggesting that simple reactions to a single cue may engage distinct cognitive processes beyond the current model.

### EEG results: event-related potentials

We focused our EEG analysis on equal choices (with additional control analysis on EEG data from forced choices), because both reward certainty and preference bias played major roles in shaping the behavioral performance of that condition. Trial-average ERPs were formed for each participant, with epochs time-locked to reward cue onset in equal and forced trials. Different reward certainty levels produced similar grand-average ERP waveforms during equal (Fig. 5A) and forced (Fig. 5B) choices, with a negative peak in the 100 – 150 ms time window (the N100 component) and a positive peak in the 300 – 400 ms time window (the P300 component).

**Figure 5.**
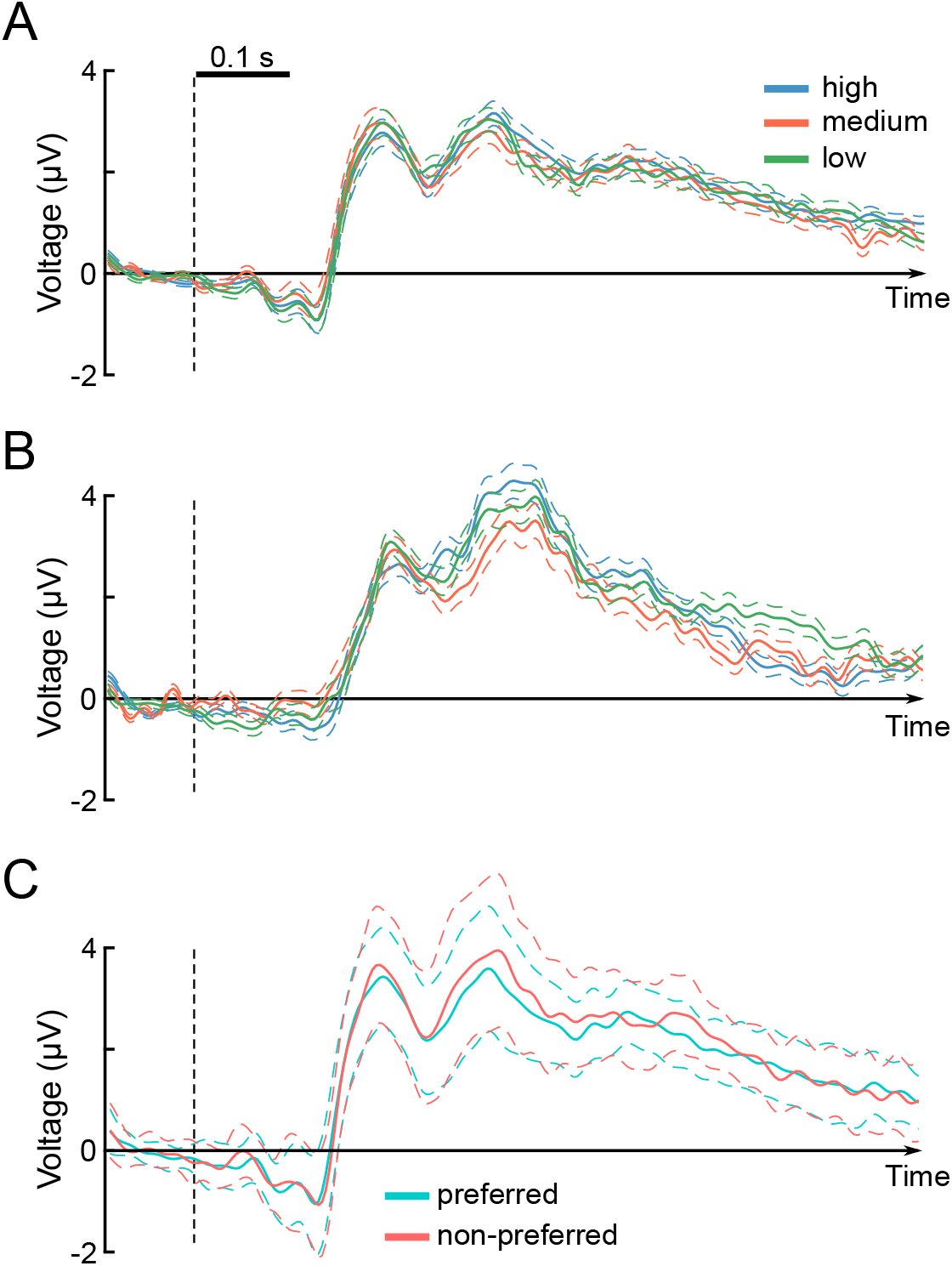
Grand-average stimulus-locked ERPs across all EEG electrodes. (**A**) ERPs from high (100%), medium (80%) and low (20%) reward certainty in equal trials. (**B**) ERPs from high (100%), medium (80%) and low (20%) reward certainty in forced trials. (**C**) ERPs from equal trials in which the preferred or non-preferred cue was chosen. In all panels, the dashed lines represent standard errors across participants.

When assessing the effect of reward certainty on ERPs, we found no univariate differences survived correction for multiple comparisons in equal (*p* > 0.552, cluster-level permutation test across 32 electrodes and all time points) or forced choices (*p* > 0.175, cluster-level permutation test). For equal trials, we found no significant difference in ERPs between preferred and non-preferred choices (Fig. 5C, *p* > 0.208, cluster-level permutation test). Therefore, in the current study, univariate ERPs were not sensitive to reward certainty or preferred/non-preferred choices.

### EEG results: multivariate patterns in equal choices

To decode multivariate information of reward certainty in equal choice trials, we applied the linear SVM on multivariate EEG patterns across all electrodes (see *Multivariate pattern analysis*). The binary classification between high and medium reward certainty was significantly above chance (*p* < 0.003, cluster permutation correction) from 120 ms after cue onset (Fig. 6A). Similarly, the information between high and low reward certainty was decodable above chance from 132 ms after cue onset (*p* < 0.042, cluster permutation correction). Relevance spatial patterns based on SVM’s weight vector (Haufe et al., 2014) showed that mid-line central and posterior electrodes contained the most information for significant classification. We found no significant classification accuracy between medium and high reward certainty (*p* > 0.073 in all time points, uncorrected).

**Figure 6.**
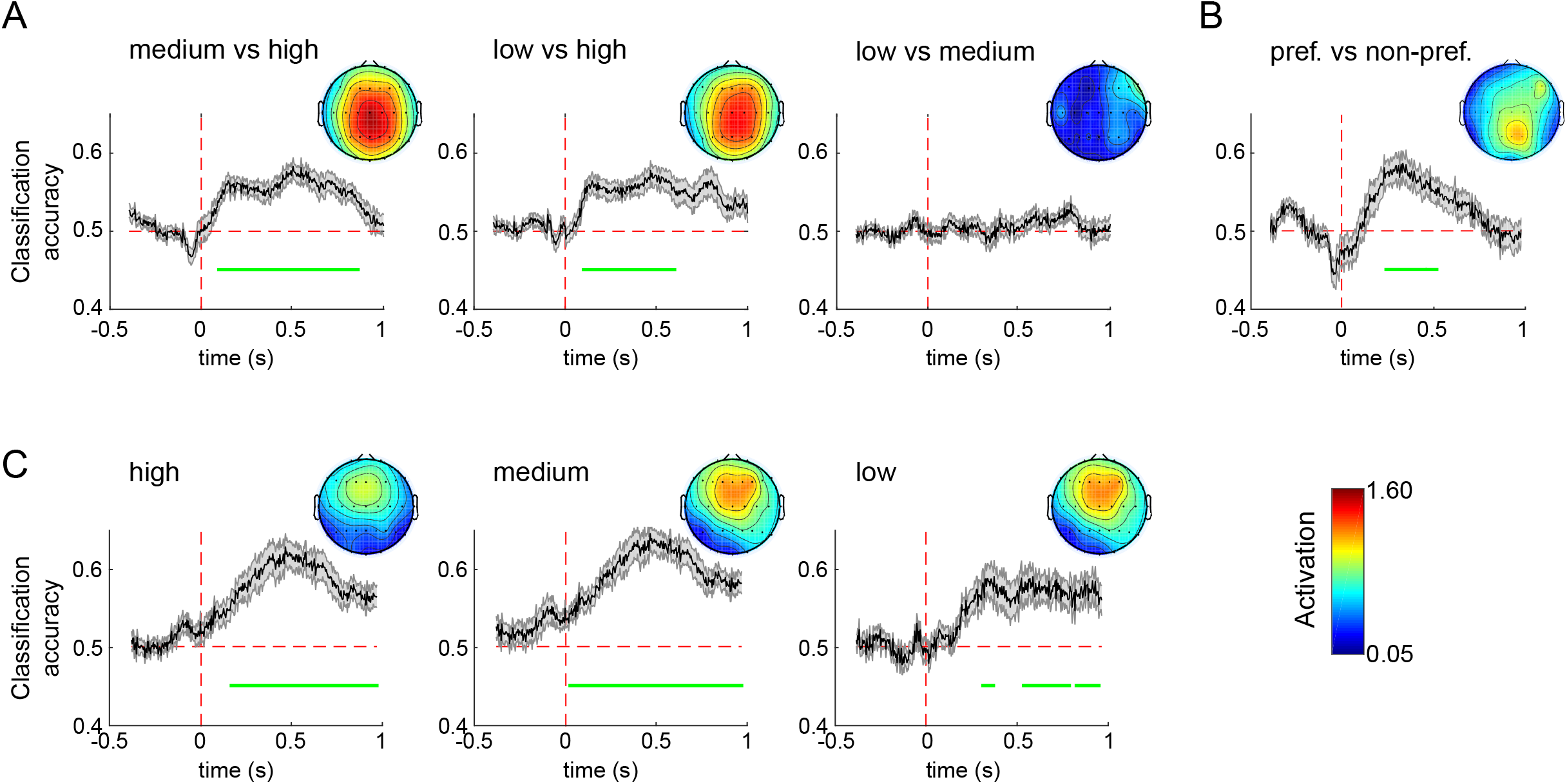
MVPA results. (**A**) Classification accuracies across time-points between equal choices with different levels of reward certainty. (**B**) Classification accuracies across time-points between equal trials with preferred and non-preferred choices. (**C**) Classification accuracies across time-points between equal and forced choices with the same level of reward certainty. In all panels, the black lines denote classification accuracies from a stratified 10-fold cross-validation and the gray areas denote standard errors. Significant decoding time windows (green horizontal bars) were determined from cluster-level permutation tests (*p* < 0.05, corrected). Topographic maps represent activation patterns from classification weights, which indicate the contribution of different EEG channels to overall classification accuracies.

We applied a similar classification procedure to discriminate the information between equal trials in which the participants chose their preferred or non-preferred choices across reward certainty levels. The information about preferred versus non-preferred choices was decodable from 212 ms to 424 ms after cue onset (*p* < 0.003, cluster permutation correction).

### EEG-informed cognitive modelling

Previous studies supported that the P300 component is associated with evidence accumulation during decision-making (Johnson, 1993; Kok, 2001; Nieuwenhuis et al., 2005; Twomey et al., 2015). Considering that the latency of early visual processing (e.g., N100 peak latency) is a part of non-decision time external to the evidence accumulation process (Nunez et al., 2018), we further hypothesized that the evidence accumulation process initiated at N100 peak latency. This led to a theoretical prediction that the slope of the rise in EEG activity between N100 and P300 peak amplitudes reflected the accumulation rate on a trial-by-trial basis. To validate this prediction, we estimated the N100 and P300 components from single trials of equal choices (Fig. 7A), using an SVD-based spatial filter to improve the signal-to-noise ratio of single-trial ERPs (see *Estimation of single-trial ERP components*). This single-trial EEG estimate was then added as a linear regressor (Equation 1) of the mean accumulation rate to the LBA model variant with the best fit to behavioral data (i.e., model 15 in Fig. 3A).

**Figure 7.**
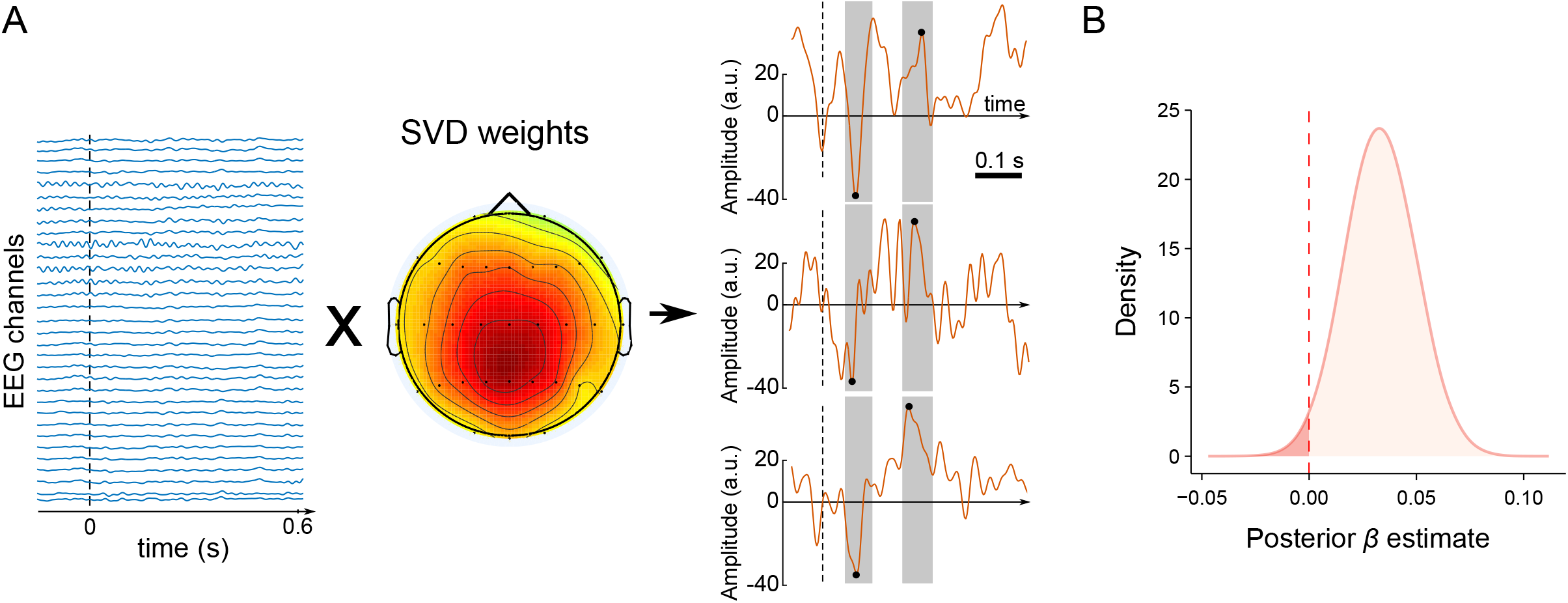
EEG-informed modelling. (**A**) The schematic diagram of extracting single-trial ERP components. 32-channel EEG signals from a single trial were multiplied by the weights of the first SVD component, calculated from the grand-averaged ERP. Next, the N100 and P300 components in that trial were identified by searching for the peak amplitude in a time of 60-164 ms for the N100 component, and 272-376 ms for the P300 component, respectively. ERP marks in three representative trials were illustrated in the right column of the panel. The ratio between N100-P300 peak amplitude difference and N100-P300 peak latency difference was calculated as a single-trial regressor for modelling. (**B**) Posterior estimates of the coefficient between the EEG-informed single-trial regressor (i.e., the rising slope of N100-P300 components) and changes in the accumulation rate.

We used the same MCMC procedure to fit the extended LBA model with the EEG-informed regressor to the behavioral data of equal choices. The extended LBA model showed good convergence (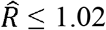 for all parameters) and provided a better fit, with a lower LOOIC score 2687 than the model without the EEG-informed regressor (LOOIC score 2796), suggesting that the rising slope of N100-P300 indeed affected the decision process. The posterior estimate of the regression coefficient *β* suggested strong evidence for a positive single-trial effect (Fig. 7B, *P_p|_*_D_ = 0.983), indicating that a bigger N100-300 slope is associated with faster accumulation rate.

### Additional analyses: alternative EEG regressors and representations of choice types

Is it possible that a simpler EEG-based regressor based on a single ERP component could provide a better model fit than the N100-P300 slope? To test this possibility, we tested four additional extended LBA models with different single-trial EEG regressors applied to the mean accumulation rate: N100 peak latency, N100 peak amplitude, P300 peak latency and P300 peak latency. All the alternative regression models showed inferior fits (LOOIC scores larger than 2700) than the one with the regressor of the N100-P300 slope. Therefore, the effects of single-trial EEG activity on the accumulation rate related to multiple ERP components.

We did not observe above-chance classification between equal trials with medium and low reward certainty (Fig. 6A). One may concern whether the lack of significant classification was due to the small number of trials in those conditions. To rule out this possibility, we conducted binary classifications to discriminate equal and forced trials with the same reward certainty. The information about trial types (equal vs. forced) was decodable at every level of reward certainty (Fig. 6C, *p <* 0.05, cluster corrected), including the one with the least number of trials (i.e., the low reward certainty). This result was expected, given the large difference in stimulus presentation and behavioral performance between the two types of choices. SVM-based relevance patterns highlighted the middle central and frontal electrodes to contain most of the information of trial types. These results suggested that the difference in classification accuracies between reward certainty levels could not be readily caused by differences in the number of trials.

## Discussion

We provided novel evidence that reward certainty and spontaneous preference shape voluntary decision processes and their electrophysiological signatures. When choosing between options with equal reward certainty, higher reward certainty facilitated RT, resulted in a *certainty effect*. We further observed a *preference effect*: biasing towards one of the two equal options with a higher choice probability and faster RT. The information of certainty and preference could be reliably decoded from multivariate ERP patterns early during decisions, but not from univariate EEG activities. Using hierarchal Bayesian implementation of a cognitive model, we showed that reward certainty and preference bias were associated with changes in the accumulation rate, a model-derived parameter to account for the speed of evidence accumulation. The accumulation rate was further affected, on a trial-by-trial basis, by the slope of the rise in ERPs between the N100 and P300 components. Together, the current study provided neurocognitive explanations for the behavioral effects during voluntary decisions.

The certainty effect on RT persisted from equal choices to simple reactions to cue locations in forced trials (Fig. 2D). In unequal choices, there was also a negative association between RT and the sum of reward certainty of the two choices (Fig. 2E). Therefore, we observed a general tendency of accelerating ones’ responses in the presence of more certain reward, even though the reward was not contingent upon the RT. These results were akin to the influence of reward magnitude in humans, which also demonstrated a facilitating effect on RT (Schurman and Belcher, 1974; Chen and Kwak, 2017). In non-human primates, the phasic activation of dopamine neurons in the ventral midbrain has similar response profiles to changes in reward certainty and magnitude (Fiorillo et al., 2003), suggesting a common mesolimbic dopaminergic pathway underlying different facets of reward processing that affect decision-making.

Bayesian model comparison identified specific effects of reward certainty on accumulation rates, highlighting two possible cognitive origins of the certainty effect. First, higher reward certainty resulted in larger mean accumulation rates in equal choices (Fig. 4), consistent with previous studies on perceptual and value-based decisions (Pirrone et al., 2014; Teodorescu et al., 2016). The accumulation rate has been linked to the allocation of attention on the task (Schmiedek et al., 2007). Because reward plays a key role in setting both voluntary (top-down) and stimulus-driven (bottom-up) attentional priority (Libera and Chelazzi, 2006; Raymond and O’Brien, 2009; Krebs et al., 2010; Yantis et al., 2012, Won and Leber, 2016), high reward certainty may boost the attentional resources allocated to sensory processing for more rapid decisions.

Second, reward certainty affected the variability of accumulation rates across trials (Fig. 4). Higher accumulation rate variability has been associated with better-memorized items (Starns and Ratcliff, 2014; Osth et al., 2017; Tillman et al., 2017). It is possible that stimuli with higher reward certainty are memorized more strongly (Miendlarzewska et al., 2016), a hypothesis to be confirmed in future studies.

Furthermore, MVPA of stimulus-locked ERPs showed multivariate EEG patterns to distinguish between certainty levels as early as 120 ms after stimulus onset (Fig 5A, see also Thomas et al., 2013), and model comparisons found no evidence to support the non-decision time to vary between certainty levels (Fig. 3A). Considering the average RT of 600-900 ms in equal choices, our results ruled out the latency of post-decision motor preparation, which constitutes a part of the non-decision time (Karahan et al., 2019), to be the main source of the certainty effect.

Interestingly, we observed a monotonic but nonlinear relationship between reward certainty and RT in equal choices: the difference between certain (100%) and uncertain (80% and 20%) reward was greater than that between the two uncertain conditions. This pointed to a special status of the 100% reward certainty distinct from lower certainty levels, as the latter always carried a non-zero risk of no reward. The salient representation of the 100% reward certainty was further highlighted by the lack of significant EEG pattern classification between the two lower certainty levels (i.e., 80% vs. 20%, Fig. 6A). Here, the certainty effect in rapid voluntary decisions resembles risk-averse behavior in economic decisions (Kahneman and Tversky, 1979), which overweighs outcomes with high reward certainty relative to less probable ones.

When choosing between equally valued options, classical evidence accumulation theories predicts a deadlock scenario with a prolonged decision process (Bogacz et al., 2006). This was not supported by recent experimental findings in value-based decisions (Pais et al., 2013; Pirrone et al., 2014), including the current study, in which equal choices took no longer than unequal ones. Our behavioral, modelling and EEG analyses indicated a preference bias, which effectively served as a cognitive mechanism to break the decision deadlock. Compared with non-preferred options, preferred decisions facilitated RTs, associated with larger accumulation rates and evoked distinct EEG multivariate patterns. The preference effect on RT was independent of reward certainty (Fig. 2A) and maintained further in forced trials (Fig. 2D), suggesting its systematical presence as a decision strategy. Because the cue-reward mapping was initially randomized and later changed within each session, this preference bias was not due to stimulus salience but established spontaneously (Voigt et al., 2019). Multiple factors may contribute to the establishment of preferred options, including simple heuristics or outcomes of early choices that unbalance subjective evaluations of equal cues (Izuma and Murayama, 2013; Bakkour et al., 2018). Future studies could validate these hypotheses by employing more frequent cue-reward re-mapping throughout experiments.

Our study highlighted the advantages of EEG-informed cognitive modelling to inform behavioral data. Hierarchical Bayesian parameter estimation of the LBA model provided a robust fit to an individual’s behavioral performance with less experimental data needed than other model-fitting methods (Vandekerckhove et al., 2011; Wiecki et al., 2013; Zhang et al., 2016). By integrating single-trial EEG regressors with the cognitive model, we identified the accumulation rate to be affected by the rate of EEG activity changes between visual N100 and P300 components. This results contributed to a growing literature of EEG markers of evidence accumulation processes, including ERP components (Twomey et al., 2015; Loughnane et al., 2016; Nunez et al., 2017), readiness potential (Lui et al., 2018) and oscillatory power (van Vugt et al., 2012). It further consolidated the validity of evidence accumulation as a common computational mechanism leading to voluntary choices of rewarding stimuli (Summerfield and Tsetsos, 2012; Maoz et al., 2019), beyond its common applications to perceptually difficult and temporally extended paradigms.

The EEG-informed modelling built upon the known functional link between the P300 component and evidence accumulation for decisions (Polich et al., 1996; Nieuwenhuis et al., 2005; Verleger et al., 2005). A new extension in the current study was to consider the accumulation process begins at the peak latency of the visual N100 component. Theoretically, the delayed initiation of the decision process accounted for information transmission time of 60∼80 ms from the retina (Schmolesky et al., 1998). Single-unit recording concurred this pre-decision delay, as neurons in putative evidence accumulation regions exhibited a transient dip and recovery activity independent of decisions, approximately 90 ms after stimulus onset (Roitman and Shadlen, 2002). Practically, our EEG data had a clear N100 component, and time-resolved MVPA identified significant pattern differences between task conditions at a similar latency. Nevertheless, we used simple stimuli with no perceptual noise. The accumulation process may start later in a trial in experiments involving more complex processing of visual information (Nunez et al., 2018). Further research could dissect the non-decision time (White et al., 2014; Tomassini et al., 2019) and compare latencies of visual encoding across decision tasks and stimuli at different levels of complexity.

Two issues require further consideration. First, our cognitive modeling was not meant to reproduce all the rich behavioral features in the data. To obtain sufficient observations for model-fitting, we combined the data before and after cue-reward re-mapping. As a result, our model did not account for behavioural changes related to cue re-mapping. Further study could employ a multi-session design to investigate how learning new cue-reward mappings influence model parameters (Zhang and Rowe, 2014). Second, we focused on the certainty and preference effects by fitting the LBA model only to the data from equal choices. Although simulations indicated that the fitted model provided similar behavioral patterns as in the empirical data in unequal and forced choices, it was not fitted directly to the experiment data in those two choice conditions. A more parsimonious model for all three types of choices would require additional assumptions, which is beyond the scope of the current study. For example, to incorporate the large RT discrepancy between equal and forced choices, one could assume that the urgency signal (Boehm et al., 2016; Thura and Cisek, 2017) plays a more dominant role in accelerating RT when no apparent comparisons are needed in forced choices.

In conclusion, when choosing between equal valued options, reward certainty and preference bias selectively modulated decision processes and their electrophysiological signatures. These findings extended and substantiated the computational framework of evidence accumulation for voluntary decisions. Our results further highlighted the intricate nature of human behavior susceptible to external factors as well as endogenous heuristics.

## Acknowledgements

This study was supported by a European Research Council starting grant (716321). WZ was supported by a PhD studentship from Cardiff University School of Psychology. DK was supported by a PhD studentship from the Engineering and Physical Sciences Research Council (1982622). We thank Sabina Baltruschat for assisting in data collection, and Craig Hedge and Petroc Sumner for comments.

